# Propulsive design principles in a multi-jet siphonophore

**DOI:** 10.1101/465245

**Authors:** Kelly R. Sutherland, Sean P. Colin, John H. Costello, Brad J. Gemmell

**Affiliations:** Oregon Institute of Marine Biology, University of Oregon, Eugene, OR 97402; Whitman Center, Marine Biological Laboratory, Woods Hole, MA 02543; Marine Biology/Environmental Sciences, Roger Williams University, Bristol, RI 02809; Biology Department, Providence College, Providence, RI 02908; Department of Integrative Biology, University of South Florida, Tampa, FL 33620

**Keywords:** propulsion, colony, velum, *Nanomia bijuga*

## Abstract

Coordination of multiple propulsors can provide performance benefits in swimming organisms. Siphonophores are marine colonial organisms that orchestrate the motion of multiple swimming zooids for effective swimming. However, the kinematics at the level of individual swimming zooids (nectophores) have not been examined in detail. We used high speed, high resolution microvideography and particle image velocimetry (PIV) of the physonect siphonophore, *Nanomia bijuga*, to study the motion of the nectophores and the associated fluid motion during jetting and refilling. The integration of nectophore and velum kinematics allow for a high-speed (maximum ~1 m s^−1^), narrow (1-2 mm) jet and rapid refill as well as a 1:1 ratio of jetting to refill time. Overall swimming performance is enhanced by velocity gradients produced in the nectophore during refill, which lead to a high pressure region that produces forward thrust. Generating thrust during both the jet and refill phases augments the distance travelled by 17% over theoretical animals, which generate thrust only during the jet phase. The details of velum kinematics and associated fluid mechanics elucidate how siphonophores effectively navigate three-dimensional space and could be applied to exit flow parameters in multijet underwater vehicles.

**Summary statement:** Colonial siphonophores produce high speed jets and generate forward thrust during refill using a flexible velum to achieve effective propulsion.

## Introduction

The physonect siphonophore *Nanomia bijuga* is a colonial cnidarian capable of long distance migrations (Robison et al., 1998) as well as short sprints and maneuvering (Costello et al., 2015). As in other physonect siphonophores, multiple swimming units, called nectophores, are organized linearly along a central nectosome (Fig. 1). The nectophores produce pulsed, high speed jets and in *N. bijuga* the nectophores are coordinated to produce forward swimming, reverse swimming and turning (Mackie, 1964; Costello et al., 2015). The coordination of multiple jets makes *N. bijuga* a highly effective swimmer that performs extensive diel vertical migrations, travelling 100s of m to the surface each night and then returning to depth during the day (Robison et al., 1998). Due to this pronounced migration behavior, its cosmopolitan distribution (Totton 1965), and the acoustic scattering properties of the gas-filled pneumatophore at the colony tip, *N. bijuga* is a prominent member of the sound scattering layer in much of the world’s oceans (Barham, 1963).

**Figure 1.**
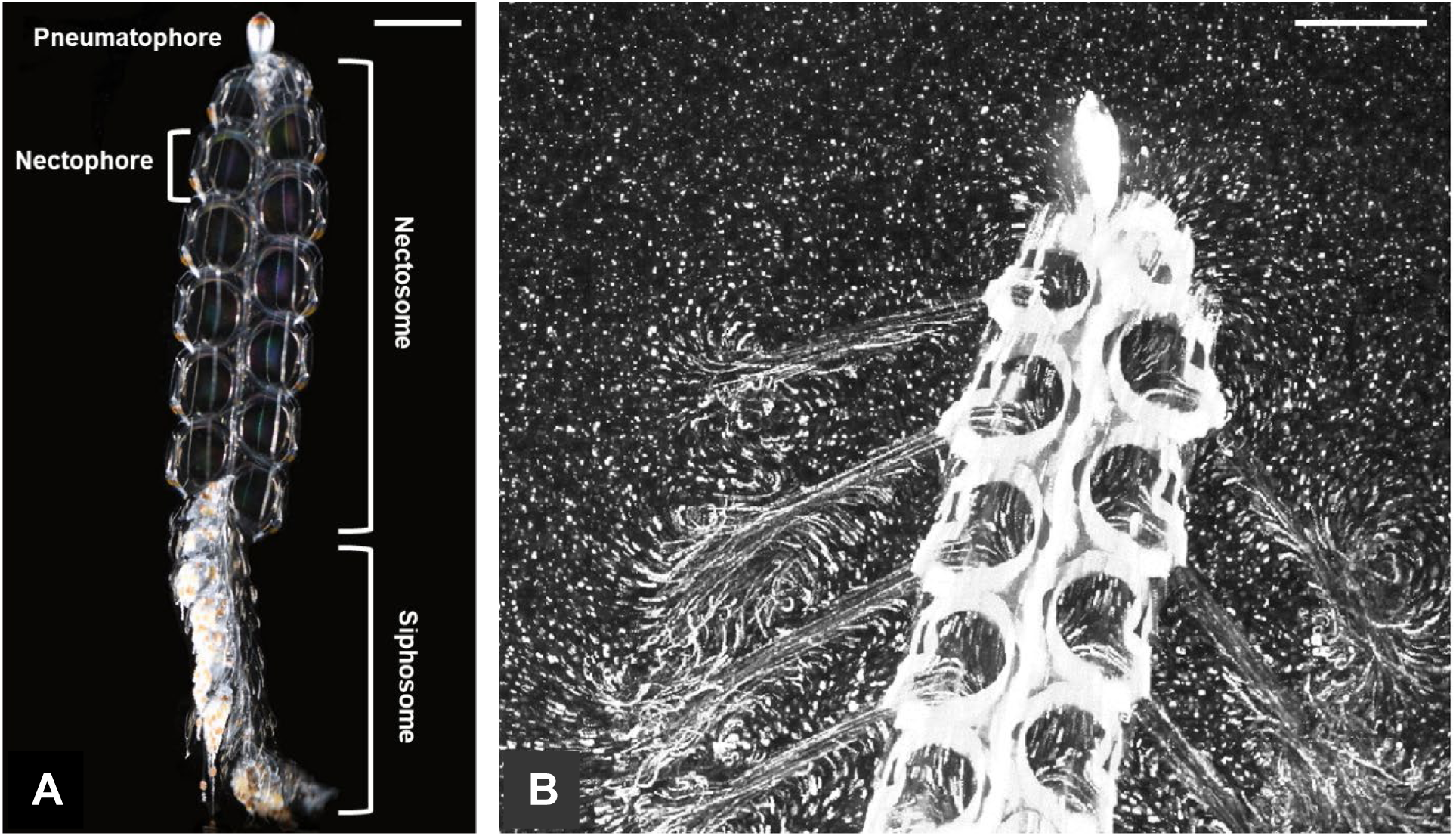
The colonial siphonophore *Nanomia bijuga* uses multiple swimming units (nectophores) (a) to produce narrow, high-speed jets (b) shown by particle tracer paths. Scale bars are 3 mm.

Up to this point, studies of swimming in *N. bijuga* have primarily focused on colony-level coordination between the nectophores, which play different roles during turning and straight swimming depending on their development and position along the nectosome (Costello et al., 2015). Newly budded nectophores toward the colony apex (Dunn and Wagner, 2006) are used for turning the colony. To achieve this, these nectophores are positioned at the apex to give a long lever arm and are oriented laterally which contributes to greater torque. Older nectophores towards the base of the colony are used for generating straight-swimming thrust. As such, they are oriented downward and produce thrust oriented more normal to the body. This coordination of nectophores allows *N. bijuga* to respond to light (Mackie, 1962) and mechanical stimuli (Mackie, 1964) and to quickly re-position between feeding bouts.

Though the division of roles among nectophores is integral in the swimming performance of *N. bijuga*, propulsion of the colony is fundamentally driven by individual-scale attributes at the level of the nectophores. However, the propulsive attributes have not been investigated in detail. Propulsion by individual cnidarian hydromedusae provides a relevant basis for comparison because the nectophores have a comparable morphology and swimming function to solitary medusa (Mackie, 1964). In particular, hydromedusae and siphonophores both possess a velum, a thin band of tissue encircling the opening of the jet orifice that reduces the jet opening and serves to manipulate the fluid flow during jetting and refill. In solitary medusae, the velum has been shown to improve swimming proficiency by allowing more fluid to roll up into the developing wake vortex, thereby increasing the jet formation number, a property associated with hydrodynamic efficiency (Dabiri et al., 2006). The velum diameter changes over the course of a jet cycle to control jet propagation (Dabiri et al., 2006). Experiments in the laboratory with changing nozzle diameters have provided further evidence to support the idea that changing the nozzle diameter leads to more effective jet formation (Allen and Naitoh, 2005; Dabiri and Gharib, 2005; Rosenfeld et al., 2009).

Just as the velum is important for directing flow in individual medusae, this structure plays a role in controlling propulsion by individual nectophores in a siphonophore colony. The variable diameter and maneuverability of the velum are likely important for directing flow into and out of individual nectophores. However, velum-associated fluid mechanics have not been investigated because of the inherent challenges of studying a fast-swimming yet fragile zooplankter. *N. bijuga* requires careful collection and handing and, once in the lab, imaging is challenging because the relevant fluid motion occurs within and near the nectophore at sub-millimeter scales but the colonies are 100s of mms in length and travel several body lengths per second.

In the present study we addressed how individual nectophore kinematics, especially of the velum, control fluid motion and contribute to effective swimming in *N. bijuga*. Understanding kinematics and fluid mechanics in detail helps resolve the mechanistic basis for effective propulsion and long-range migration in *N. bijuga*. We show that the motion of the nectophore and velum are carefully coordinated to produce narrow high speed jets, effective for proficient forward swimming coupled with rapid refill over a much larger velar-opening area with lower velocities, which minimizes rearward thrust and hydrodynamic loss. A comprehensive understanding of propulsive principles in this multi-jet biological swimmer may have application to underwater vehicles that rely on single or multiple jets.

## Methods

*Nanomia bijuga* colonies were hand-collected from docks at Friday Harbor Labs in May and June of 2014, 2016 and 2017. Colonies were maintained in running seawater tables at ambient environmental temperatures (10-12 °C). *N. bijuga* were gently transferred to glass vessels prior to filming and observations were made within 24 hrs. of collection. Glass vessels ranged in size from 8×2×4 to 15×4×12 (height × depth × width in cm) to accommodate colonies of varying sizes. All kinematic and fluid mechanics measurements were taken from colonies that were swimming upward.

### Kinematics

For kinematic measurements of individual nectophores, *N. bijuga* colonies were illuminated using a brightfield collimated light set-up with a 10× LWD objective lens (Gemmell et al., 2014). Images were collected with high speed monochrome video cameras at 500-6,400 frames per s. and resolutions of 1024×1024, 1280×1024, or 2048×2048 pixels, depending upon the frame rate necessary to temporally resolve the specific process. Side view and plan views of the nectophore were taken during forward swimming. Image stacks were imported into the open-source software program ImageJ to measure the change of the velum opening over at least one complete pulse cycle. For larger field of view images of whole-colony swimming, the same brightfield set-up was used with a 50 mm lens. Time spent jetting and refilling during each pulse cycle was extracted from large field of view videos.

### Fluid mechanics

To examine the relationship between velum kinematics and fluid mechanics of the jet, we used a combination of brightfield particle image velocimetry (PIV) (Gemmell et al., 2014), laser sheet PIV (Sutherland et al., 2014) and particle tracking (Costello et al., 2015). The brightfield PIV set up was identical to the set up for kinematics but the water was seeded with *Isochrysis* cells (Size= 5-6 μm). For the laser sheet PIV set-up, 10 μm hollow glass beads were seeded into the tank and illuminated with a <1 mm thick laser light sheet (532 nm). Image sequences during jetting and refill were selected where the velar aperture was bisected by the laser sheet. Image pairs were subsequently analyzed in DaVis 8.3.1 (LaVision GmbH, Goettingen, GER) using a cross-correlation PIV algorithm with a decreasing interrogation window size of 64 × 64 pixels to 32 × 32 pixels or 32 × 32 pixels to 16 × 16 pixels with 50% overlap to produce velocity vectors and vorticity contours. Instantaneous jet velocities were typically too high to capture with PIV even at shutter speeds of 1/80,000 s. Therefore, jet velocities were made over several pulse cycles by tracking individual particles from image stacks in ImageJ at an increment of 0.05 s (Costello et al., 2015). Instantaneous nectophore area and velar diameter were measured from the same image sequences to directly compare kinematics and resultant fluid velocities. To verify that the fluid velocities measured from particle tracks were reasonable, the volume flux (*Q* = *A_j_ vt*) during jetting and refill were compared based on the mean jet area (*A_j_* = *πab* where *a* and *b* are the major and minor axes of an ellipse), mean jet velocity, *v*, and time, *t* (of jet or refill).

### Swimming performance calculation

Swimming speed, *U*, of free-swimming *N. bijuga* over time, *t*, during jetting and refill were calculated from image sequences based on the change in x, y position of the top of the nectosome just below the pneumatophore as:

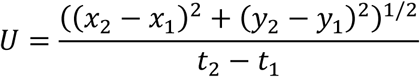

To assess whether refill kinematics and fluid kinematics impart performance benefits, colony velocity during refill was compared to the velocity of a passive model travelling based on inertia alone. The distance travelled during refill for both *N. bijuga* (n=5) and the passive model were calculated based on the respective velocities.

The post-contraction velocity of the passive projectile model was estimated following the equations of ballistic motion (e.g. Gemmell et al., 2013). The initial post-contraction velocity was assumed to be the maximum velocity of the *N. bijuga* colony. Since live colonies were swimming vertically, we ignored the horizontal component and calculated projectile motion from a balance of forces in the vertical direction

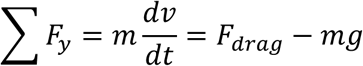

The model was assumed to be a smooth cylinder with same diameter and height as the nectosome. This is a conservative estimate since the colony trailing structures were not included in the length and these structures add drag, which would further decelerate the colony. Based on the parameters of the nectosome, equation 2 becomes

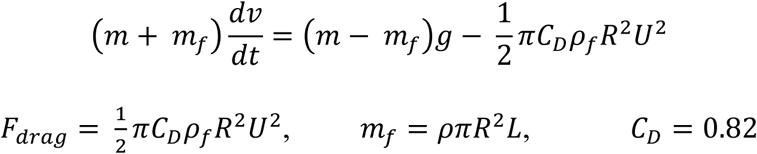

where *m_f_* is the added mass of the fluid around a cylinder, *U* is the velocity of the siphonophore, *R* is the radius, *m* is the mass of the siphonophore, *ρ_f_* is the density of seawater and *C_D_* is the drag coefficient for an elongate cylinder. We used the fourth-order Runge–Kutta method for computing 2D motion of a body through a fluid, which is detailed by Biringen and Chow (2011), to obtain instantaneous position and velocities. A time step Δt of 0.01 s was used, and siphonophores were assigned a *ρ* of 1,025 kg m^3^ because they are slightly negatively buoyant relative to the water (*ρ* of 1,020 kg m^−3^).

### Pressure field

Pressure field data in and around the nectophore were calculated from the measured velocity vectors as previously described (Colin et al., 2012; Dabiri et al., 2014) using the Dabiri et al., 2014 queen2 MATLAB algorithm (dabirilab.com/software). High resolution video (2048×2048 pixels) at 1000 frames s^−1^. was required to achieve sufficient information for pressure calculations inside the nectophore. The queen2 algorithm calculates pressure from velocity by numerically integrating the inviscid Navier-Stokes equation. To reduce the effects of errors from individual paths, a median-polling scheme is used to integrate multiple paths to calculate an estimate of pressure at every point in space (Dabiri et al., 2014). Applications of the algorithm to study other swimming organisms (Gemmell et al., 2015; Lucas et al., 2017) provided validation of the methods.

## Results

The motion of the velum was highly dynamic during jetting and refill (Fig. 2). A plan view of the velum during a representative jetting/refill cycle shows that while relaxed, the velum was triangular in shape. During forward jetting, the velum aperture constricted to become smaller and more circular in shape. During refill, the velum opened rapidly (Fig. 2A, Video S1). A time-series showing a side view of the velum during a jetting/refill cycle showed that the velum moved laterally and curved downward during jetting to position the jet more parallel to the body. During refill, the velum reversed direction and moved inside the nectophore to accommodate fluid filling the nectophore (Fig. 2B, Video S2).

**Figure 2.**
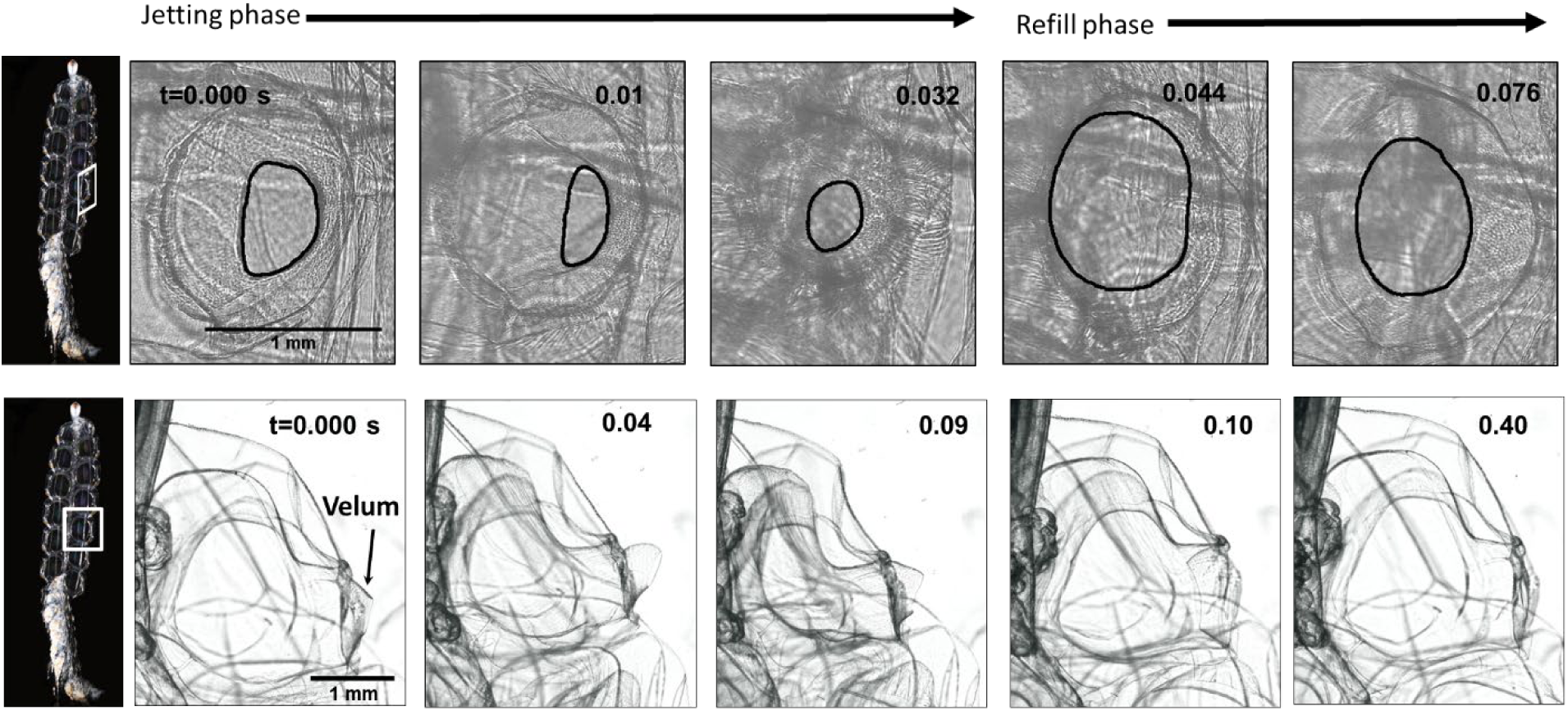
Plan and side view of velum and nectophore during forward swimming showing change in velum diameter and shape. Also see videos S1, S2.

The rapid contraction of the velum during jetting (Fig. 2) drove a narrow high speed jet (Fig. 3). During refill, velocities were an order of magnitude lower and fluid was pulled in perpendicular to the colony orientation and also upstream of the velum. None of the fluid appeared to be pulled from downstream of the velum (Fig. 3).

**Figure 3.**
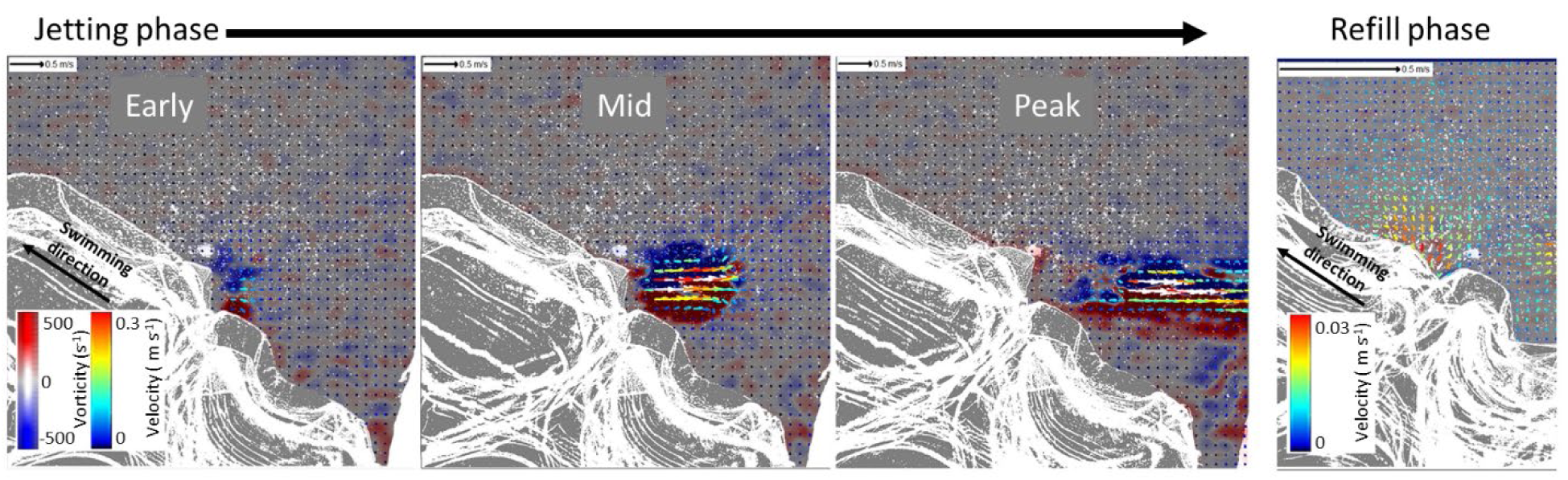
Side view of nectophore showing velocity vectors and vorticity contours during jetting and refill. Image height=4 mm. Also see video S3.

The relative timing of the velum closing and nectophore contraction controlled the jet properties (Fig. 4). During jetting, the velum closed first and then nectophore contracted 0.05-0.08 s later so that the nectophore aperture began closing before fluid was ejected (Fig. 4A). The resultant jet reached peak speeds of ~1 m s^−1^ (mean ± SD=899±183 mm s^−1^, n=4; Figs 3, 4) with vorticity reaching ~500 s^−1^ (Fig. 3). Peak jet velocities were associated with periods when the velum diameter was smallest (Fig. 4B) and the greatest rate of change in jet velocity was associated with the greatest rate of change in nectophore diameter (Fig. S1). During refill, the velum relaxed and nectophore expanded simultaneously but the rate of opening was faster in the velum (Fig. 4A). Refill velocities slowed to a minimum velocity that was an order of magnitude lower than the jetting velocity (Figs 3, 4) and the fluid velocity was relatively steady with minimal acceleration (Fig. 4B). The amount of volume flux during jetting (mean ± SD=17.1±3.3 mm^3^) was not significantly different from volume flux during refill (mean ± SD=16.4±1.9 mm^3^) (t-test: t=0.36, p=0.76, n=3), which helped confirm that velocity measurements were accurately measured.

**Figure 4.**
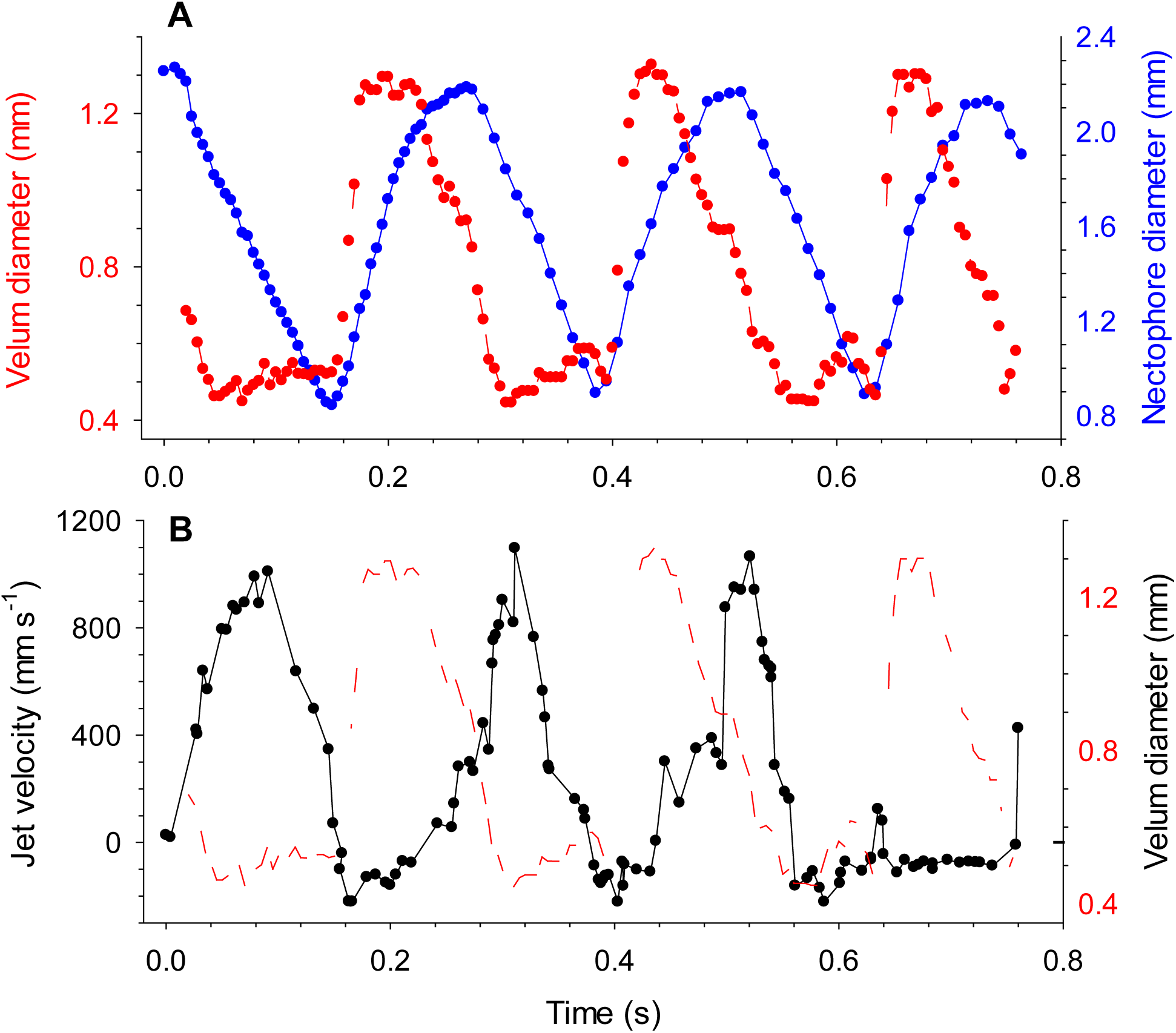
*N. bijuga* nectophore kinematics (a) showing change in velum and nectophore diameter over three pulse cycles and (b) the resultant jet velocities plotted with change in velum diameter.

As a result of the rapid movements of the velum and nectophore, the time spent jetting (mean ± SD= 0.122± 0.025 s (57.9%)) and refilling (mean ± SD=0.106± 0.068 s (42.1%)) were not significantly different (~1:1 ratio; Fig. 5, paired t-test, t=0.73, p=0.49, n=9). However, the time spent refilling was more variable than the time spent jetting (Fig. 5). On average a full jet cycle was completed in 0.23 ± 0. 077 s.

**Figure 5.**
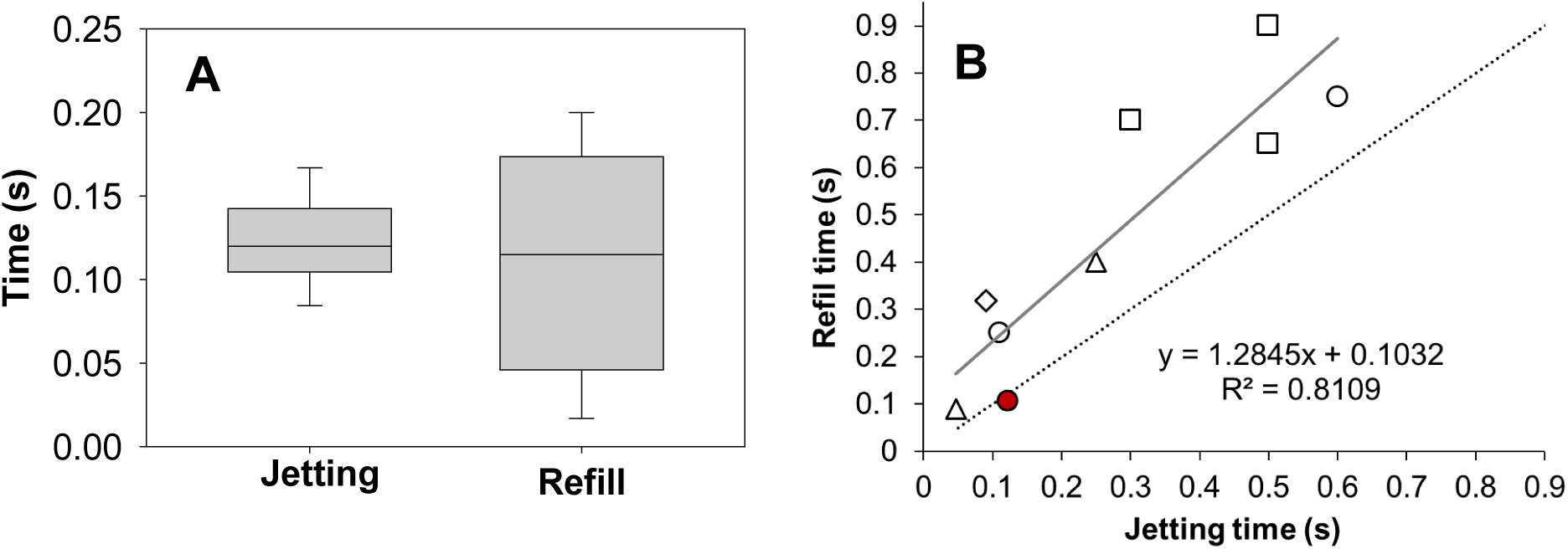
Jetting versus refill time. (A)Time spent jetting (mean± SD= 0.122± 0.025 s) and refilling (mean± SD= 0.106± 0.068 s) during forward swimming by *N. bijuga* (n=9). (B) Jetting time plotted with refill time for jet propelled swimmers between 0.1 and 2 cm in length. Siphonophores (○) with *N. bijuga* in red (This study, Bone and Trueman 1982), pelagic tunicates (Δ) (Bone and Trueman 1983, 1984), larval squid (◊) (Andersen and Demont 2000), hydromedusae (□) (Gladfelter 1972, Colin and Costello 2002). Grey line is regression fitted to all data points and dotted line represents 1:1.

The orchestrated kinematics and fluid mechanics during swimming resulted in a measureable performance benefit for *N. bijuga*. During refill, the colony continued to move forward (Fig. 6). Though the colony decelerated during refill, the rate of deceleration was slower than would be expected for a passive model with the same dimensions and density as *N. bijuga*. As a result, *N. bijuga* traveled significantly further— by 17%— during each pulse cycle than would be expected based on a passive model (Fig. 6). A quantification of the fluid mechanics and associated pressure fields during refill helps to explain the mechanism underlying this performance benefit. The morphology of the nectophore allowed fluid to be pulled in laterally from water upstream of the nectophore opening. The fluid was accelerated as it moved through the velar opening and along the wall of the nectophore (Fig. 7A). As the fluid was forced towards the inner wall of the nectophore it was also deflected upwards and as it continued to move along the wall of the nectophore it decelerated so that when the fluid reached the entrance of the nectophore again it had decelerated considerably. This movement of fluid against the inner walls resulted in temporary vortex being formed within the nectophore during each refill. The pressure fields within the nectophore (Fig. 7B), calculated from the velocity vectors suggest that the performance benefit experienced by *N. bijuga* was due to advantageously located high pressure fields. Fluid movement within the nectophore itself resulted in fluid being forced against the inner and upper walls of the nectophore, which in turn generated high pressure in these regions of the nectophore. This has the potential of generating upward thrust against the nectophore wall due to the lateral orientation of the nectophore, which produces lateral thrust instead of downward thrust as water enters during refilling. A negative pressure region created by the rotating inside the nectophore would have no direct influence on thrust because the negative pressure region does not interact with any surface of the nectophore (Fig. 8). The net result of the pressure field is therefore an upward thrust, which helps to explain the performance benefit experienced by *N. bijuga* during refill (Fig. 6).

**Figure 6.**
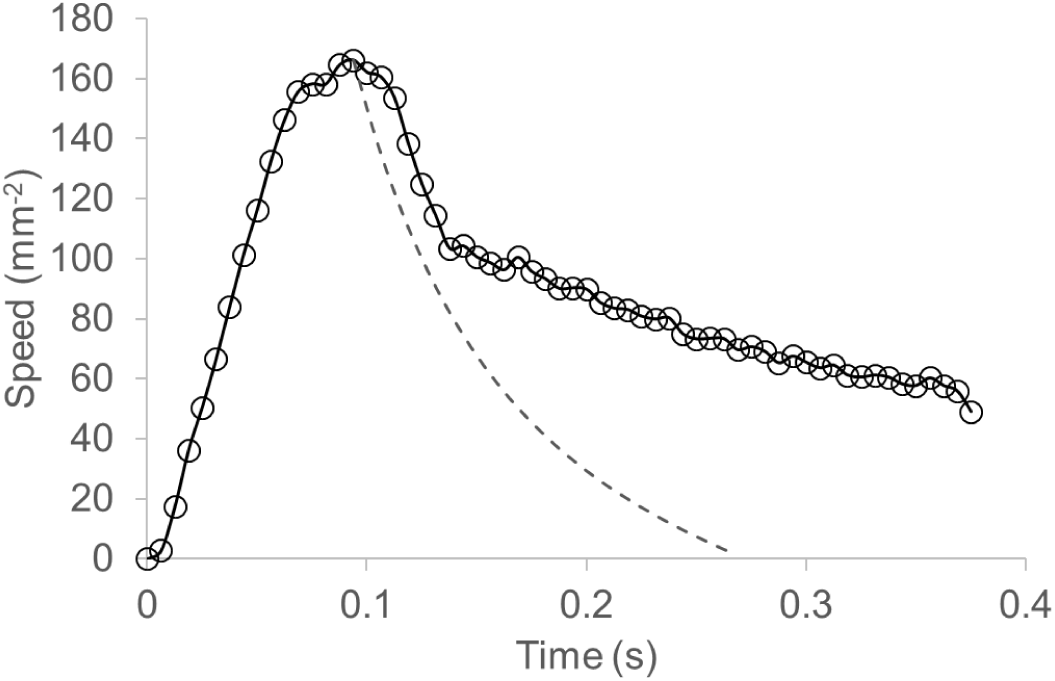
Performance benefits during refill. Representative profile showing *N. bijuga* colony speed over one full jet/refill cycle. Dashed line shows expected speed of a model based on inertia alone.

**Figure 7.**
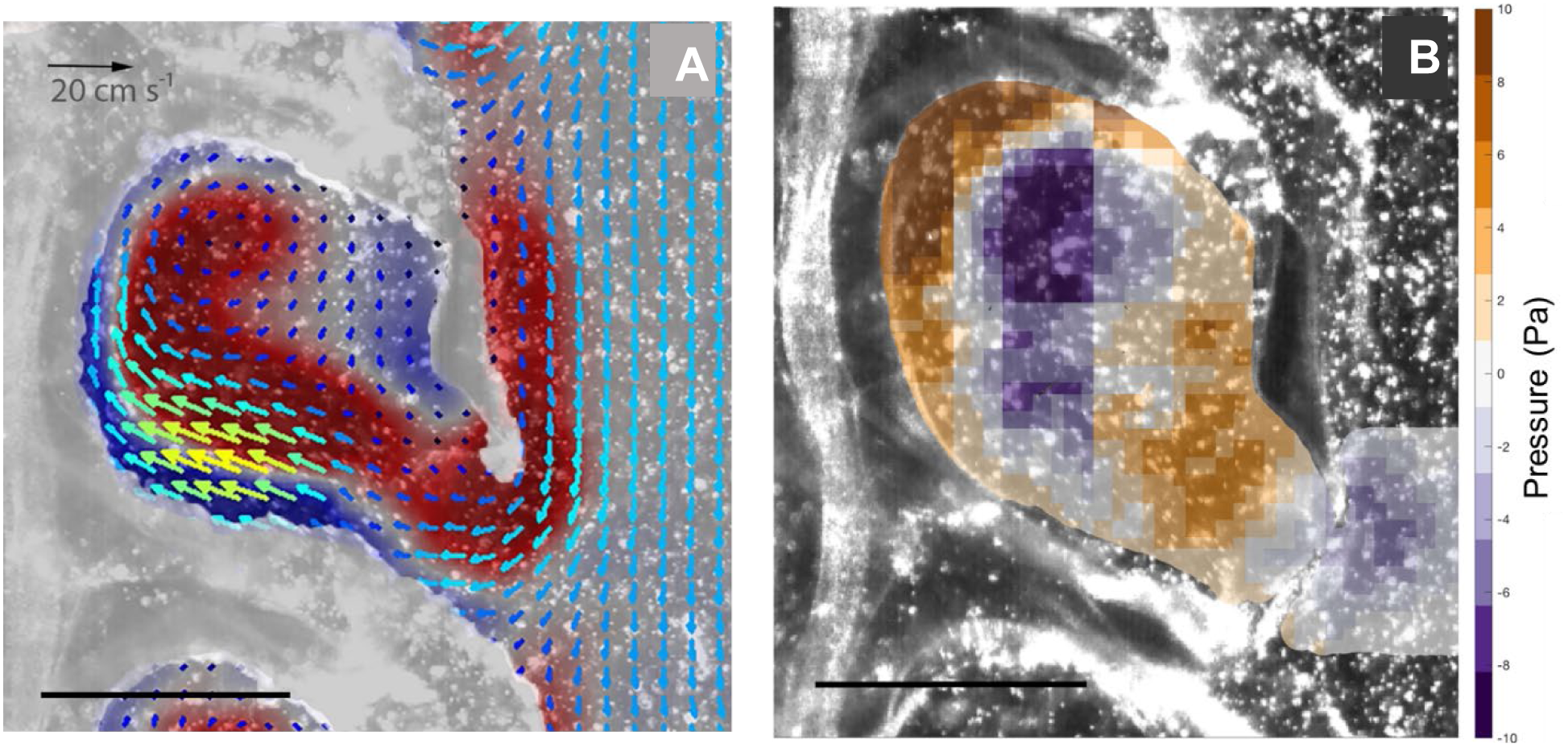
Fluid mechanics during nectophore refill. (A) Velocity vectors and vorticity contours and (B) pressure contours during refill by a single nectophore. Scale bars are 3 mm. Also see Video S4.

**Figure 8.**
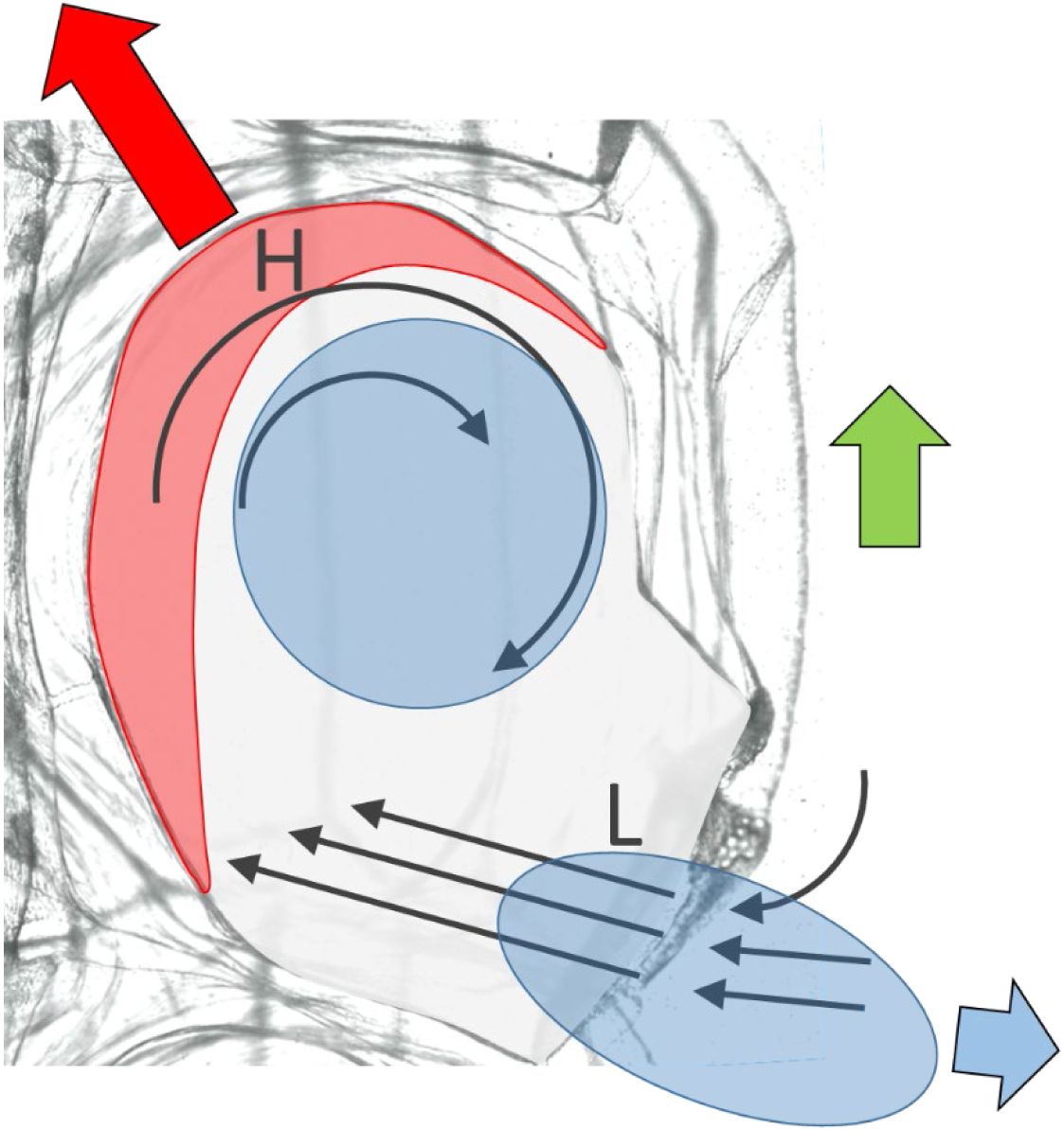
Proposed model detailing fluid flow and corresponding pressure fields during refill responsible for generating performance benefit in *N. bijuga*. Velocity vectors (thin arrows), high (H) and low (L) pressure and resultant thrust vectors (wide arrows) are shown.

## Discussion

Here we show that at the individual nectophore scale, a highly maneuverable velum and tight coordination between the kinematics of the nectophore and velum produce impressive jet speeds and rapid refill between jets. The details of kinematics and fluid mechanics at the level of individual nectophores have been elusive because *N. bijuga* is fragile and requires careful collection. Additionally, the nectophores produce small scale yet high velocity jets over short time intervals (Fig. 3), which requires high shutter speeds and a sufficiently large field of view with high resolution. We find that the velum is highly maneuverable and changes diameter and shape rapidly to control fluid during jetting and refill (Fig. 2). Orchestration of the velum diameter and nectophore diameter drive a high speed narrow jet and a low speed refill (Fig. 4). The velum closes slightly in advance of the nectophore contraction so that as the volume of the nectophore decreases the velum opening also decreases allowing for a narrow high velocity jet. The velum then opens very rapidly (~0.03 s) and the large area of the velar opening slows the fluid down as it is pulled inside the nectophore. Fluid is pulled primarily from upstream of the nectophore potentially mitigating rearward thrust that would be generated if fluid was drawn in from downstream of the nectophore (Figs 3, 7). In addition to directing the fluid, the velum’s rapid movements result in a 1.15 ratio of time spent jetting to time spent refilling and a short overall jet cycle time (0.23 s) compared to other jet propelled swimmers (Fig. 5).

Jetting in cnidarian medusae has previously been characterized as asymmetric with a shorter time spent jetting relative to refill time (Costello et al., 2008) and this pattern holds across most other jetting marine organisms. Ratios of jetting to refill are 0.28 in the paralarval squid, *Doryteuthis pealeii* (Anderson and Demont, 2000), between 0.43 and 0.77 in hydromedusae (Gladfelter, 1972; Colin and Costello 2002), 0.63 in the salp, *Salpa fusiformis* (Bone and Trueman, 1983) and 0.52 in *Doliolum* spp. (Bone and Trueman, 1984). Previously, the highest reported jetting to refill ratio was for another siphonophore, *Abylopsis tetragona* (Bone and Trueman, 1982) (Fig. 5B). This difference in jetting to refill time has been attributed to aspects of kinematics and fluid mechanics. In hydromedusae, contraction of the bell during jetting is governed by subumbrellar muscles. During refill, the muscles relax and the energy stored in the elastic mesoglea of the bell returns the bell to a relaxed position. The short contraction time results in higher fluid velocities during jetting than during relaxation/refill because the volume moves out of the bell aperture over a shorter time period during contraction compared to refill. The lower refill velocity may serve to decrease rearward thrust, which would impede the animal’s forward motion.

In *N. bijuga* the refill time closely matches the jetting time but the potentially negative consequences in terms of high refill velocities that might create rearward thrust are mitigated by a highly variable velum diameter (Fig. 2) that slows fluid intake considerably during refill (Figs 3, 4). In other words, the velum increases the aperture area very rapidly during refill to slow the intake velocity. Because volume is conserved, there are two variables that control fluid velocity during jetting or refill: orifice opening area and time. *N. bijuga* appears to be exceptional in controlling the orifice area allowing for a reduced refill time. The ratio of jetting to refill is not completely static, however. The relative timing of jetting versus refill is variable for *N. bijuga* (Figs 2, 5). In other jet propelled swimmers it has been remarked that swimming speed typically increases by decreasing refill time but maintaining the jetting time (Hydromedusa: Gladfelter, 1972; Squid: Andersen and Demont, 2000). During high speed sprints that might occur during an escape, avoiding predation likely outweighs hydrodynamic losses due high velocity refill.

This study reveals that the presence of a variable aperture—the velum—is a key feature in controlling swimming wake dynamics. During a jetting cycle, the velum in *N. bijuga* undergoes dynamic three-dimensional changes in shape, size and orientation. While all hydromedusae possess a velum there is a large variability among species in how pronounced the velum is relative to the bell diameter. Species that have a more developed velum are more proficient at swimming and typically characterized as ambush predators (Colin and Costello 2002). In the fast-swimming hydromedusa *Nemopsis bachei*, flow visualizations showed that time-varying velar kinematics enhance the vortex structures produced during jetting to maximize thrust (Dabiri et al., 2006). A variable velar opening appears to be a critical feature in enhancing wake structures of fast-swimming hydromedusae that have been quantified.

In addition to changing diameter, the velum of *N. bijuga* also changes shape. During jetting, the velum goes from a two-dimension shelf to a three-dimensional truncated cone shape. During rest, the opening shape is triangular and then as the velum contracts, the opening shifts to an ellipse and finally a circular opening (Fig. 2). Other jetters with variable exit openings also exhibit shape changes. The shape of a squid’s siphon goes from a circular shape to an ellipse (Anderson and Demont, 2000) and in salps, the exit nozzle goes from a circular shape to a curved ellipse (Sutherland and Madin, 2010). A triangular shaped exit nozzle has not been observed in other jet-propelled organisms but, interestingly, the aortic valve in the mammalian heart is triangular in shape (Yoganathan et al., 2004). Leonardo da Vinci was fascinated by this triangular shape and associated fluid mechanics and considered the human aortic valve to be an example of a highly efficient pump (Robiseck, 1991). Fluid dynamics associated with changes in exit diameter of jet nozzles have been investigated in biological swimmers (Dabiri et al., 2008) and with jet nozzles in the lab (Allen and Naitoh, 2005; Dabiri and Gharib, 2005; Rosenfeld et al., 2009); however, the importance of change in shape of the nozzle opening remains an open area of investigation.

The jetting phase is typically the focus of studies of aquatic jet propulsion but recent work has underscored the importance of the refill phase in garnering performance benefits (Gemmell et al., 2013; Gemmell et al., 2018). In cnidarian medusae the stopping vortex that moves into the bell during refill generates pressure gradients to produce passive energy recapture (PER). In *Aurelia aurita*, PER generates a 30% increase in the distance travelled during each swimming cycle and contributes to the low cost of transport for cnidarian medusa (Gemmell et al., 2013). While PER is an important feature across species of cnidarian medusae, the degree of performance benefit varies among species with different swimming modes ranging from 0 to a 60% increase in total distance travelled (Gemmell et al., 2018).

*N. bijuga* travels 17% farther during refill than would be expected for a passive model; i.e. *N. bijuga* that does not produce thrust during refill (Fig. 6). This benefit—the equivalent of shaving 40 min off a 4 hr marathon time—leverages the fluid dynamics associated with the refill period. In addition to a low refill velocity and pulling fluid upstream of the nectophore as discussed above (Figs 3, 4), the circulation of fluid in the nectophore produces regions of high and low pressure (Fig. 7). As the fluid enters the nectophore, it accelerates and is forced into the walls to produce a broad high pressure region toward the upper section (proximal and medial) of the nectophore. This high pressure region would therefore be expected to produce a thrust vector with a substantial upward component that would contribute to forward swimming (Fig. 8). In contrast, the sub-ambient pressure at the entrance to the velum is relatively low and therefore produces a smaller thrust vector. Further, because the fluid is drawn in at an oblique angle (5-35°) relative to the colony axis (Costello et al., 2015), this thrust vector would be expected to have only a small downward component. The upward thrust generated by high pressure at the upper wall and reduction of the downward thrust through the lateral orientation of the nectophore opening creates predominantly lateral instead of downward thrust and thus explains the performance benefit during the refilling phase. This mechanism can only function when multiple jetting units are present as the lateral force created during refill in one nectophore is offset by the equal and opposite force produced by the nectophore on the opposite side.

Swimming capabilities can be important for migrating, feeding and escaping predation. For some cnidarian hydromedusae, swimming and feeding are tightly coupled such that swimming also generates vortices that pass through the tentacles bringing prey into contact with these feeding structures (Costello and Colin, 2002). In jetting hydromedusae, which tend to have more streamlined prolate forms, swimming is decoupled from feeding. These hydromedusae are characterized as ambush predators; they spend much of their time motionless (Colin et al., 2003) and rely on swimming prey intercepting the tentacles. In colonial siphonophores, the swimming structures are completely decoupled from the feeding structures both spatially and functionally.

The nectophores, which are in a linear array along the nectosome, have a specific swimming role and the feeding structures are situated further down the colony along the siphosome (Fig. 1A).

Placed in an adaptive context, the decoupling of swimming and feeding functions may have allowed for additional performance benefits in swimming for colonial siphonophores (Mackie, 1986; Costello et al., 2008). Nectophores are arranged in a streamlined linear architecture and have an exclusive propulsive role that is coordinated by nerves. The colony can therefore perform diverse swimming movements, much like a well-integrated solitary organism. Propulsion by a single apical nectophore produces turning and forward swimming can be achieved by synchronous or asynchronous pulsing of the nectophores (Costello et al., 2015). Asynchronous swimming is slower than synchronous swimming but more hydrodynamically efficient because the colony swims at a steady speed (Sutherland and Weihs, 2017). An additional benefit of coloniality is that colonies are less size-constrained than individual medusae. The loosening of allometric size constraints, decoupling of swimming from feeding at the individual nectophore scale and coordination among nectophores helps to explain why many siphonophore species are proficient swimmers (Bone and Trueman, 1982; Costello et al., 2015) capable of long-distance diel vertical migrations (e.g. Pugh 1984; Andersen et al., 1992).

## Acknowledgements

We thank Friday Harbor Laboratories for facilities’ support to conduct this research.

---

**Figure S1.**
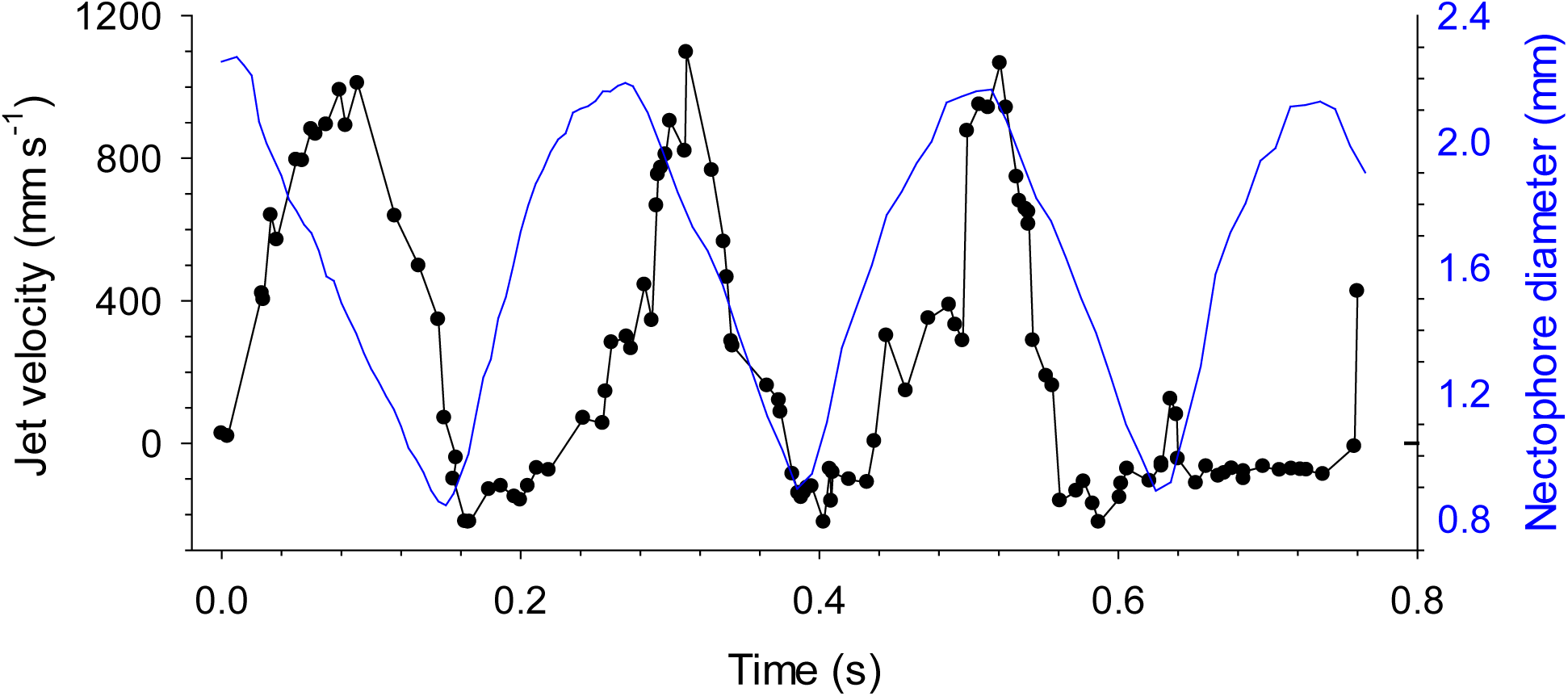
*N. bijuga* jet velocity over three pulse cycles plotted with change in nectophore diameter.

## References

Allen, J. J. and Naitoh, T. (2005). Experimental study of the production of vortex rings using a variable diameter orifice. Phys. Fluids. 17, 061701.

Andersen, V., Sardou, J. and Nival, P. (1992). The diel migrations and vertical distributions of zooplankton and micronekton in the Northwestern Mediterranean Sea. 2. Siphonophores, hydromedusae and pyrosomids. J. Plankton Res. 14, 1155-69.

Anderson, E. J. and Demont, M. E. (2000). The mechanics of locomotion in the squid *Loligo pealei*: locomotory function and unsteady hydrodynamics of the jet and intramantle pressure. J. Exp. Biol. 203, 2851-2863.

Anderson, E. J. and Grosenbaugh, M. A. (2005). Jet flow in steadily swimming adult squid. J. Exp. Biol. 208, 1125-1146.

Barham, E.G. (1963). Siphonophores and the deep scattering layer. Science. 140, 826-828.

Biringen, S., and Chow, C. Y. (2011). An introduction to computational fluid mechanics by example. John Wiley & Sons.

Bone, Q. and Trueman, E. R. (1982). Jet propulsion of the calycophoran siphonophores *Chelophyes* and *Abylopsis*. J. Mar. Biol. Assoc. U. K. 62, 263-276.

Bone, Q. and Trueman, E. R. (1983). Jet propulsion in salps (Tunicata: Thaliacea). J. Zool. Lond. 201, 481-506.

Bone, Q. and Trueman, E. R. (1984). Jet propulsion in Doliolum (Tunicata: Thaliacea). J. Exp. Mar. Biol. Ecol. 76, 105-118.

Colin, S. P. and Costello, J. H. (2002). Morphology, swimming performance and propulsive mode of six co-occurring hydromedusae. J. Exp. Biol. 205, 427-437.

Colin, S.P., Costello, J.H. and Klos, E. (2003). In situ swimming and feeding behavior of eight co-occurring hydromedusae. Mar. Ecol. Prog. Ser. 253, 305-309.

Costello, J. H. and Colin, S. P. (2002). Prey resource use by coexistent hydromedusae from Friday Harbor, Washington. Limnol. Oceanogr. 47, 934-942.

Costello, J. H., Colin, S. P. and Dabiri, J. O. (2008). Medusan morphospace: phylogenetic constraints, biomechanical solutions, and ecological consequences. Invert. Biol. 127, 265-290.

Costello, J. H., Colin, S. P., Gemmell, B. J., Dabiri, J. O. and Sutherland K. R. (2015). Multi-jet propulsion organized by clonal development in a colonial siphonophore. Nat. Commun. 6, 8158.

Dabiri, J. O., Bose, S., Gemmell, B. J., Colin, S. P., and Costello, J. H. (2014). An algorithm to estimate unsteady and quasi-steady pressure fields from velocity field measurements. J. Exp. Biol. 217, 331–336.

Dabiri, J. O., and Gharib, M. (2005). Starting flow through nozzles with temporally variable exit diameter. J. Fluid Mech. 538, 111–136.

Dabiri, J. O., Colin, S. P. and Costello, J. H. (2006). Fast-swimming hydromedusae exploit velar kinematics to form an optimal vortex wake. J. Exp. Biol. 209, 2025-2033.

Dunn, C. W. and Wagner, G. P. (2006). The evolution of colony-level development in the Siphonophora (Cnidaria: Hydrozoa). Dev. Genes Evol. 216, 743-54.

Gemmell, B. J., Costello, J. H., Colin, S. P., Stewart, C. J., Dabiri, J. O., Tafti, D. and Priya, S. (2013). Passive energy recapture in jellyfish contributes to propulsive advantage over other metazoans. Proc. Natl Acad. Sci. USA 110, 17904-17909.

Gemmell, B.J., Jiang, H. and Buskey, E.J. (2014). A new approach to micro-scale particle image velocimetry (μPIV) for quantifying flows around free-swimming zooplankton. J. Plankton Res. 36, 1396-1401.

Gemmell, B. J., Colin, S. P., Costello, J. H. and Dabiri, J. O. (2015). Suction-based propulsion as a basis for efficient animal swimming. Nat. Commun. 6, 8790.

Gemmell, B.J., Colin, S.P. and Costello, J.H. (2018). Widespread utilization of passive energy recapture in swimming medusae J. Exp. Biol. 221, p.jeb168575.

Gladfelter, W. B. (1972). Structure and function of the locomotory system of *Polyorchis montereyensis* (Cnidaria, Hydrozoa). Helgoländer wiss. Meeresunters. 23, 38-79.

Lucas, K. N., Dabiri, J. O. and Lauder, G. V. (2017). A pressure-based force and torque prediction technique for the study of fish-like swimming. PloS one. 12, e0189225.

Mackie, G.O., (1962). Pigment effector cells in a cnidarian. Science, 137,689-690.

Mackie, G. O. (1964). Analysis of locomotion in a siphonophore colony. Proc. R Soc. B 159, 366–391.

Mackie, G. O. (1986). From aggregates to integrates: physiological aspects of modularity in colonial animals. Phil. Trans. R. Soc. Lond. B. 313, 175-196.

Pugh, P. R. (1984). The diel migrations and distributions within a mesopelagic community in the north east Atlantic. 7. Siphonophores. Prog. Oceanogr. 13, 461-89.

Robicsek, F. (1991). Leonardo da Vinci and the sinuses of Valsalva. Ann. Thorac. Surg. 52, 328-35.

Robison, B. H., Reisenbichler, K. R., Sherlock, R. E., Silguero, J. M. B. and Chavez, F. P. (1998). Seasonal abundance of the siphonophore, *Nanomia bijuga*, in Monterey Bay. Deep-Sea Res. 45, 1741–1751.

Rosenfeld, M., Katija, K., and Dabiri, J. O. (2009). Circulation Generation and Vortex Ring Formation by Conic Nozzles. J. Fluids Eng. 13, 091204.

Sutherland, K. R. and Madin, L. P. (2010). Comparative jet wake structure and swimming performance of salps. J. Exp. Biol. 213, 2967–2975.

Sutherland, K.R., Costello, J.H., Colin, S.P. and Dabiri, J.O., (2014). Ambient fluid motions influence swimming and feeding by the ctenophore *Mnemiopsis leidyi*. J. Plankton Res. 36, 1310-1322.

Sutherland, K. R. and Weihs, D. (2017). Hydrodynamic advantages of swimming by salp chains. J. Royal Soc. Interface. 14, 20170298.

Totton, A. K. and Bargmann, H. E. (1965). A synopsis of the Siphonophora. British Museum (Natural History).

Yoganathan, A. P., He, Z. and Casey Jones S. (2004). Fluid mechanics of heart valves. Annu. Rev. Biomed. Eng. 6, 331-362.

